# *En bloc* preparation of *Drosophila* brains enables high-throughput FIB-SEM connectomics

**DOI:** 10.1101/855130

**Authors:** Zhiyuan Lu, C. Shan Xu, Kenneth J. Hayworth, Patricia Rivlin, Stephen M. Plaza, Louis Scheffer, Gerald M. Rubin, Harald F. Hess, Ian A. Meinertzhagen

**Affiliations:** Department of Psychology and Neuroscience, Life Sciences Centre, Dalhousie University, Halifax, Nova Scotia, Canada B3H 4R2; Janelia Research Campus, Howard Hughes Medical Institute, 19700 Helix Drive, Ashburn VA 20147, USA

## Abstract

Deriving the detailed synaptic connections of the entire nervous system has been a long term but unrealized goal of the nascent field of connectomics. For *Drosophila*, in particular, three sample preparation problems must be solved before the requisite imaging and analysis can even begin. The first is dissecting the brain, connectives, and ventral nerve cord (roughly comparable to the brain, neck, and spinal cord of vertebrates) as a single contiguous unit. Second is fixing and staining the resulting specimen, too large for previous techniques such as flash freezing, so as to permit the necessary automated segmentation of neuron membranes. Finally the contrast must be sufficient to support synapse detection at imaging speeds that enable the entire connectome to be collected. To address these issues, we report three major novel methods to dissect, fix, dehydrate and stain this tiny but complex nervous system in its entirety; together they enable us to uncover a Focused Ion-Beam Scanning Electron Microscopy (FIB-SEM) connectome of the entire *Drosophila* brain. They reliably recover fixed neurons as round profiles with darkly stained synapses, suitable for machine segmentation and automatic synapse detection, for which only minimal human intervention is required. Our advanced procedures use: a custom-made jig to microdissect both regions of the central nervous system, dorsal and ventral, with their connectives; fixation and Durcupan embedment, followed by a special hot-knife slicing protocol to reduce the brain to dimensions suited to FIB; contrast enhancement by heavy metals; together with a progressive lowering of temperature protocol for dehydration. Collectively these optimize the brain’s morphological preservation, imaging it at a usual resolution of 8nm per voxel while simultaneously speeding the formerly slow rate of FIB-SEM. With these methods we could recently obtain a FIB-SEM image stack of the *Drosophila* brain eight times faster than hitherto, at approximately the same rate as, but without the requirement to cut, nor imperfections in, EM serial sections.

---

Increasingly rapid progress is being made to secure the exact synaptic wiring diagram of a brain, its connectome (Lichtman and Sanes, 2008), complete at the electron microscope (EM) level. That knowledge will enable functional analyses of synaptic circuits, and so help reveal the mechanism of identified behaviors. Attention is directed mostly to the model brains of genetically manipulable species (Luo et al., 2008), especially those of the mouse and the fruit fly *Drosophila melanogaster* (Fig. 1). Neuron numbers, about 100,000 in *Drosophila* (Shimada et al., 2005; Meinertzhagen, 2018), 1,000 times fewer than in the mouse, has enabled significant progress on the former, despite the wide range of methods available for brains of other sizes (e.g. Hayworth et al., 2014; Kubota et al., 2019). However, *Drosophila*’s brain presents a special problem because even though the z-axis resolution for serial-section EM (ssEM) may be satisfactory for mouse brains (Denk and Horstmann, 2004; Hayworth et al., 2014), and while the tiny neurites of *Drosophila* neurons are shorter than those in the mouse, favoring their three-dimensional reconstruction, their caliber (typically ≤ 0.2µm) is finer, making comprehensive reconstruction in the z-axis problematic using ssEM.

**Figure 1.**
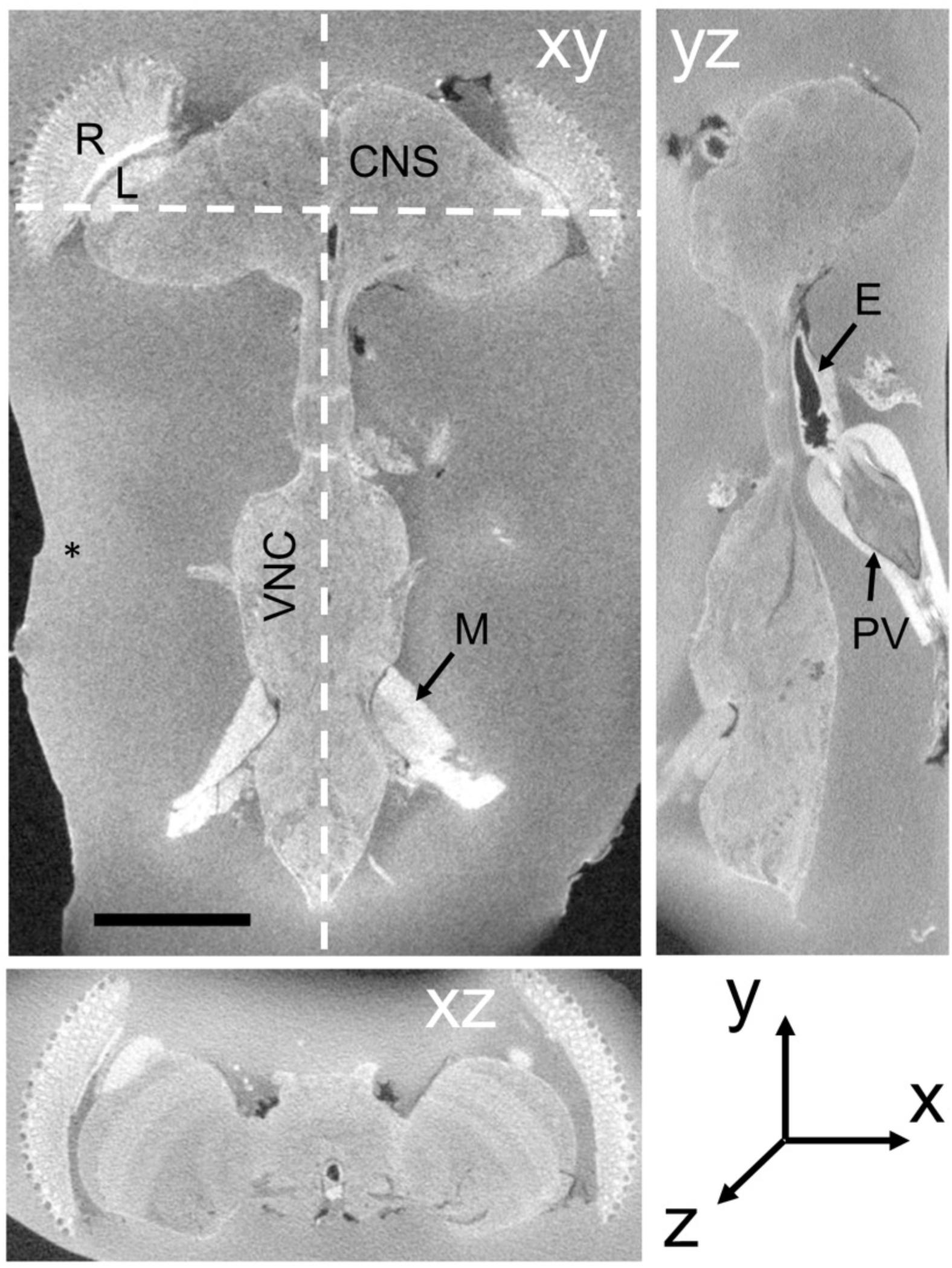
X-ray micro-CT of dissected *Drosophila* CNS in three orthogonal planes, xy, yz and xz. xy: frontal slice of the entire CNS, including the VNC. yz saggital plane. xz transverse plane. Different zones in the surrounding BSA reveal successive additions of the cross-linked protein. E: esophagus; L: lamina; M: muscle; R: retina; PV: proventriculus. Scale bar in xy: 200µm.

Overcoming these problems, FIB-SEM (Knott et al., 2008; Xu et al., 2017) is the preferred method to image *Drosophila* neuropile. Not only does it circumvent the supreme technical skill required to cut extended series of ultrathin sections for serial-section EM (SSEM), but also z-axis resolution is not limited by section thickness. An additional advantage is that z-axis resolution can be adjusted to equal that in x and y (typically 8nm for FIB-SEM for x,y and z) compared with TEM (4 nm in x, y and >40nm in z: Zheng et al., 2018) (Fig. 2). FIB-SEM thus provides the means to collect isotropic 8nm image stacks well suited to reconstruct the slender neurites of *Drosophila* (Takemura et al., 2015; Xu et al., 2017; Shinomiya et al., 2019). Providing an ideal approach to that task, this method has been adopted at the Janelia Research Campus of HHMI in an intensive effort to derive the entire connectome of a fly’s brain, one that can be comprehensively mined for circuit information (e.g. Takemura et al., 2017; Horne et al., 2018).

**Figure 2.**
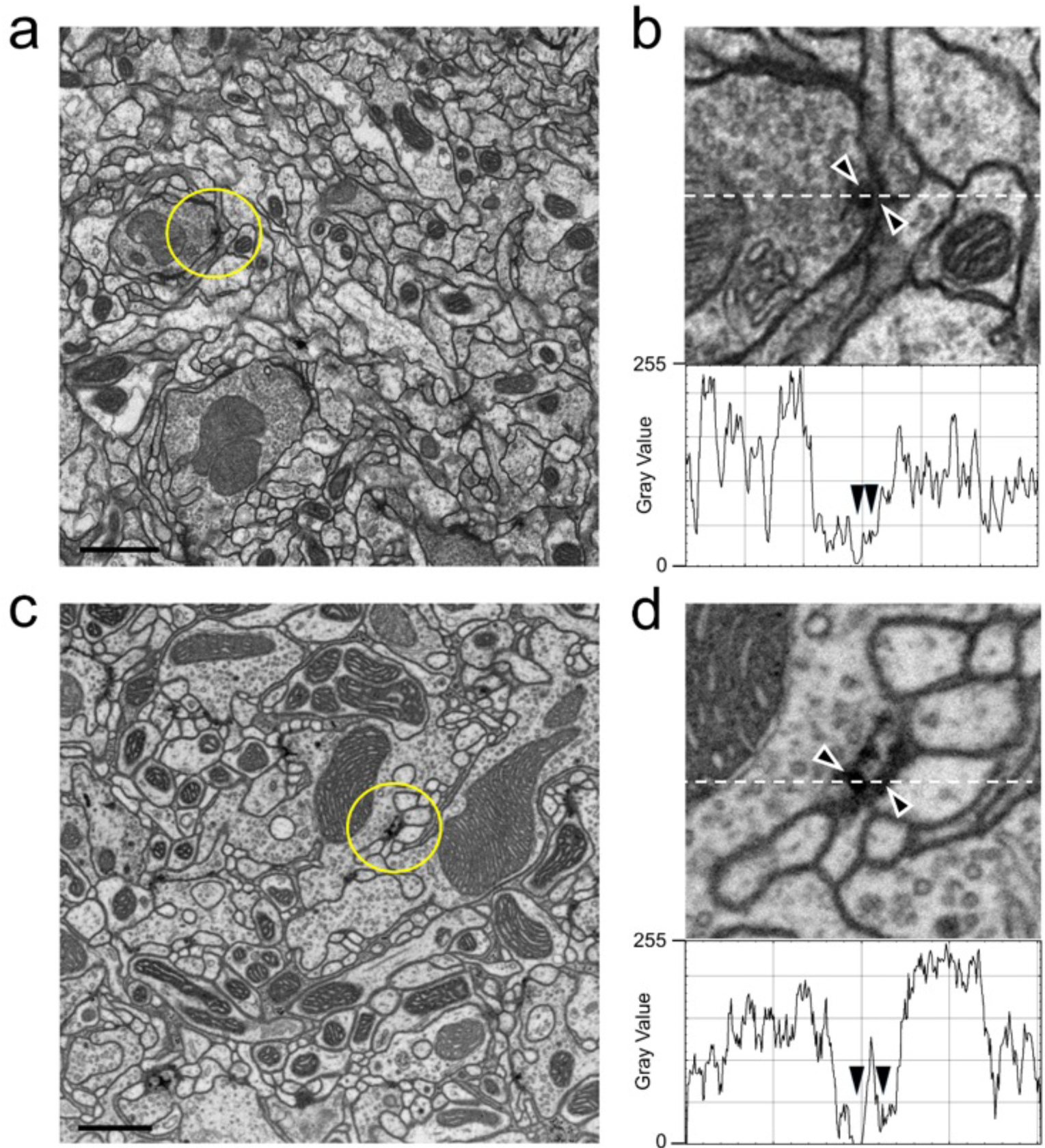
TEM (a,b) and FIB-SEM (c,d) images compared. (a) TEM image from series of 50-nm sections post-stained from the mushroom body calyx reported in Butcher et al. (2012); image resolution is 3.7nm per pixel. (b) Enlargement from (a) to show the ultrastructure of a synapse and membranes. Image is scanned in the lower panel to show the gray-scale values for staining, especially for the synapse and synaptic membranes (arrowheads). (c) Compare image quality with high-resolution 4-nm per pixel FIB-SEM image of the protocerebral bridge (see also Supplementary Video 2). (d) Representative individual synaptic profile and synaptic membranes from (c) are indicated by arrowheads in (d), pointed to the T-bar presynaptic ribbon and synaptic membranes. The image is scanned along the interrupted white line to show the grey scale value through organelles. Arrowheads for scan line indicate the electron density of the selected synapse and its membranes. Note that the contrasts of the synaptic T-bar and membrane density in the FIB-SEM image (d) both match those from TEM (b). Scale bars (a,c) 1µm.

EM resolution is required to see synaptic organelles, and the methods for fixing and staining brain tissue in *Drosophila* are well established (e.g. Meinertzhagen and O’Neil, 1991; Yasuyama et al., 2002; Prokop, 2006; Schürmann, 2016); but these have changed little in 50 years, and moreover are not well suited to FIB-SEM imaging. Here, we report various methods that we have developed within the last decade to fix and stain the sub- and supraesophageal regions of the *Drosophila* brain (Fig. 1). Together these regulate the segmental ganglia of the ventral nerve cord (VNC), the conduit for much of the brain’s biological output, motor behavior (Niven et al., 2008). Our methods are adapted to automate the segmentation of neurons in both ganglia and VNC, to identify the synaptic profiles between such neurons, and especially to increase FIB-SEM’s formerly slow rate of imaging.

## Results

We present a consolidated method for FIB-SEM of the *Drosophila* brain, based on a number of protocols (Table 1), each compiled from multiple parametric repetitions. Together with earlier methods (Fig. 2a), these have taken a decade to develop and perfect. Central to them were the development of new microdissection protocols (Fig. 3, Supplementary Fig. 1), improvements in the heavy metal staining of the brain that support faster rates of FIB-SEM imaging, and the exact targeting of specific regions using X-ray tomography of embedded stained brains.

**Figure 3.**
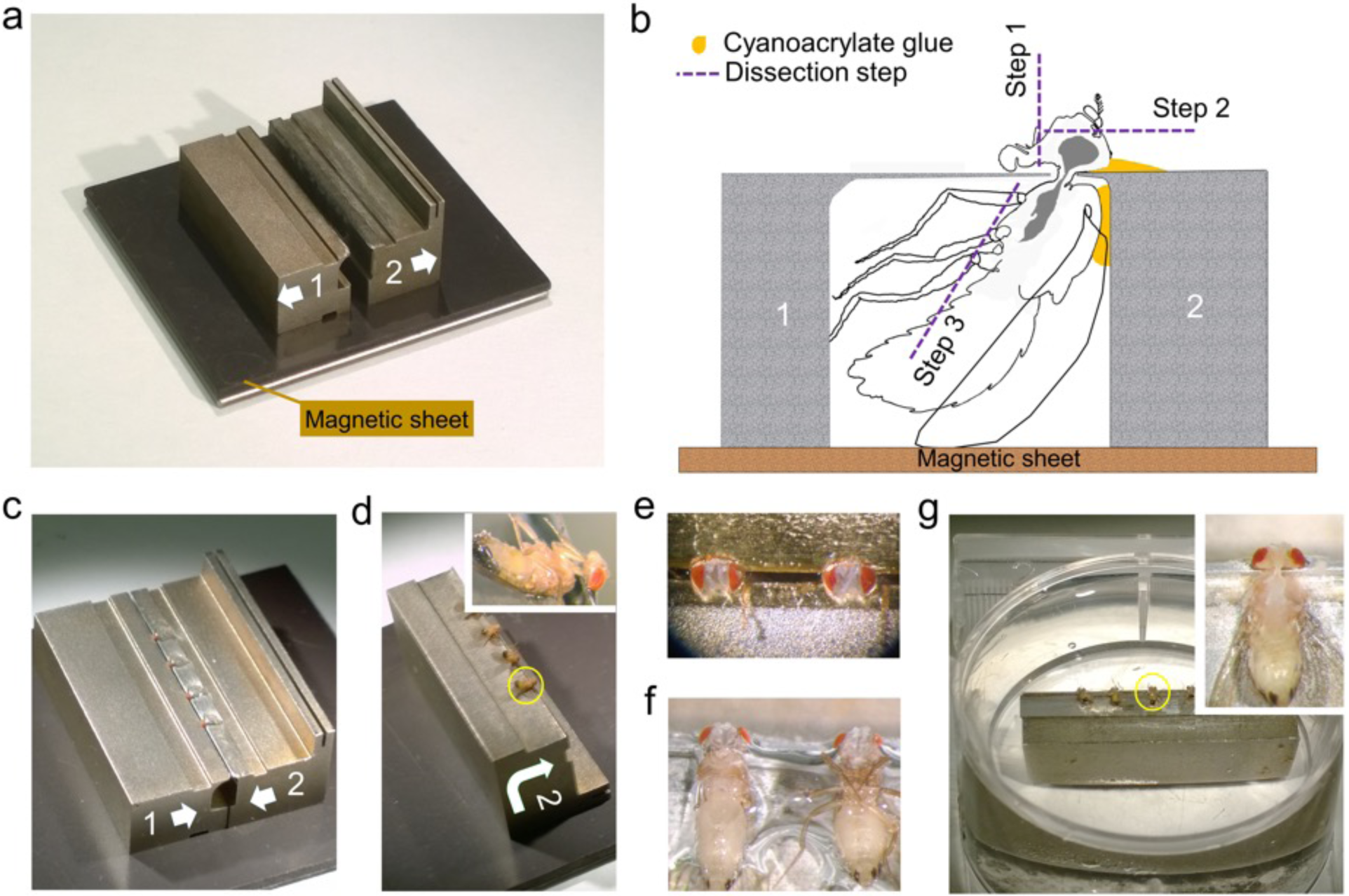
Dissection collar modified for dissecting the entire CNS (see Supplementary Video 1). **a**. Two half stocks (1 and 2) arranged to form a pillory on a magnetic base for quick assembly. Quick release shown as arrows on each stock (also in c and d). Stock 2 has an elevated bar for forceps to grip and transfer the stock. **b**. Outline of fly in a dissection pillory, held captive at its neck by thin metal shim, to show the position of Loctite glue and dissection steps 1-3. **c**: Four flies with protruding heads, loaded and glued into assembled pillory as in (b). **d**: Stock 2 separated from its partner stock 1 and turned over, with flies glued by the cuticle of their dorsal thorax as in (b). Inset: Enlarged view of one fly. **e**: Two heads in saline after dissection step 1 and 2 in (b) to remove the proboscis and frontal head cuticle together with antennae. **f**: Showing the pinioned flies turned over from (e) and transferred to saline, with left-side legs removed as in step 3 in (b). **g**: The pinioned flies on stock 2 in saline or PB buffer in tissue culture dish. Inset: Single fly with legs and abdominal cuticle removed to expose the entire CNS (brain and VNC) as shown in the yellow circle in (g).

**Table 1.** *Drosophila* sample preparation procedures and results

| Procedure and Results | Methods |  |  |  |
| --- | --- | --- | --- | --- |
|  | 1. HPF-FS | 2. PLT-LTS | 3. PLT-LTS heavy metal enhancement | 4. PLT-LTS progressive heavy metal enhancement |
| Tissue dissection | 200µm Vibratome slices | Brain tissue dissect out with metal collar | CNS dissect out with metal collar | Brain dissect out with/out metal collar |
| Pre-fixation | 2.5%GA + 2.5% PFA, RT, 15min | 2.5%GA + 2.5% PFA, RT, 2h | 2.5%GA + 2.5% PFA, RT, 2h | 2.5%GA + 2.5% PFA, RT, 2h |
| Post-fixation | HPF cryo-fixation | 0.5% OsO<sub>4</sub>, 30min, 4°C, W | 0.5% OsO<sub>4</sub>, 40min, 4°C | 1% OsO<sub>4</sub>, 40min, 4°C |
| Heavy metal enhancement |  | 0.5% UA, 30min, 0°C, W<br>0.8% OsO<sub>4</sub> 20min 0°C | no wash, change to 0.8% K ferrocyanide, 2h, 0°C + 0.5% UA, 30min, 0°C; W<br>lead aspartate, 4°C overnight; or 4h RT W<br>0.8% OsO<sub>4</sub>, 20min, 0°C | no wash, change to 1.5% K ferrocyanide, 1.5h, 0°C + 30min RT; W<br>1% Tch, 15min, RT; W, 2% OsO<sub>4</sub>, 30min, RT; W<br>0.5% UA, 30min, RT, W; lead aspartate, 30min, 55°C, & 1h, RT, W |
| *en bloc* staining and dehydration | FS:<br>95% acetone, 1% OsO<sub>4</sub>, 0.2% UA, 1% methanol, 38h - 90°C<br>14h -20°C | PLT:<br>acetone, 0°C to -25°C<br>LTS:<br>97% acetone with 1% OsO<sub>4</sub> and 0.2% UA, 30h, -25°C | PLT:<br>ethanol, 0°C to -25°C<br>LTS:<br>97% ethanol with 1% OsO<sub>4</sub> and 0.2% UA, 30h, -25°C | PLT:<br>ethanol 0°C to -25°C<br>LTS:<br>97% acetone with 0.2% UA, or 96% ethanol with 1% PTA 30h, -25°C |
| Infiltration | Acetone | Propylene oxide | Propylene oxide | Acetone |
| Embedding | Durcupan | Poly/Bed 812 | Poly/Bed 812 | Durcupan |
| Summary |  |  |  |  |
| Hot knife cutting | Not suitable | 20 µm slices | 25 µm slices on BSA coating tissue | Not suitable |
| FIB-SEM 8nm imaging scan rate | 1.25 MHz | 2 MHz | 3 MHz | 10 MHz |
| FIB-SEM 8nm imaging speed | 25x10<sup>3</sup> µm<sup>3</sup> per day per machine | 30x10<sup>3</sup> µm<sup>3</sup> per day per machine | 50x10<sup>3</sup> µm<sup>3</sup> per day per machine | 200x10<sup>3</sup> µm<sup>3</sup> per day per machine |
| FIB-SEM data collection | Medulla, lobula, lobula plate; α-lobe; antennal lobe | Hemi-brain (central complex, mushroom body and more) | Central nervous system (CNS) | Medulla<br>1st instar larva CNS |

Notes: LTS = Low temperature staining; W = washing (3 X 10 min)
Hot knife cutting properties improved by infiltration in propylene oxide not acetone, and Epon (Poly/Bed 812, Ted pella) instead of Durcupan. Hot knife introduces a ~30% increase in overall SEM imaging work. FIB-SEM data collected by Method 1 were performed in a modified Zeiss NVision40 system, while those collected by Methods 2 to 4 were performed in modified Zeiss Merlin systems. FIB-SEM 8nm imaging scan rates are calculated based on the modified Merlin platform for direct comparison.

### General features of FIB-SEM images

The general features of FIB-SEM images obtained using the updated fixation and staining method we present below are first authenticated against conventional images obtained with TEM. To make this comparison valid, we first needed to capture FIB-SEM images at higher resolution (4nm/pixel in x,y,z, Fig. 2c,d) than we routinely used (8nm/pixel) to be more nearly comparable to the TEM image of the same brain region, for which we illustrate a region of the protocerebral bridge (Fig. 2c,d). Cell and organelle profiles visible from TEM are all immediately recognizable in the FIB-SEM image, and indistinguishable using either imaging method at the magnifications chosen. Importantly for segmentation, cell membranes and synaptic profiles are all clearly visible. Pre- and postsynaptic elements were both more electron-dense than with conventional TEM methods using countertop post-staining (compare Figs. 2b, 2d). The FIB-SEM illustrated, 4nm/pixel in x,y,z, (Fig. 2c), has a higher resolution than that (8nm) at which neurons were routinely segmented for segmentation, however, each reconstructed voxel thus having a volume 2^3^ = 8 times larger. Synaptic sites (Fig. 2b,d) could be clearly detected semi-automatically from their increased electron density (Huang et al., 2018), with typically a single T-shaped presynaptic density (or T-bar), opposite which sit a number of postsynaptic processes.

### Microdissecting the fly’s brain

To prepare adult *Drosophila* brains for imaging, various previously reported dissection methods (Meinertzhagen, 1996; Wolff, 2011) are mostly too rudimentary. To visualize neurons that arborize not only in the dorsal supraesophageal brain but also in the subesophageal ganglion and VNC, we imaged each part in parallel to reconstruct both arbors of single neurons. For this, we developed a method to microdissect and fix the two ganglia of a single brain intact, together with their corresponding cervical connectives. This required that we dissect the *Drosophila* brain by holding the head in a custom machined metal collar (Heisenberg and Böhl, 1979) and then transfer the ensemble to primary fixative. The yield of well-preserved brains is not high and successfully increased only by means of such collars. About four heads each held in a single collar are together transferred to primary fixative within about 5 minutes. In most reports in the literature, the lamina is simply torn off, so that the brain’s outer margin is the medulla cortex, but in our improved methods we use careful dissection to retain the lamina, which offers its own merits as a model neuropile (Meinertzhagen and O’Neil, 1991; Meinertzhagen and Sorra, 2001). In its current application, the method is further modified into a two-stage dissection that enables us to preserve intact the supraesophageal brain together with the subesophageal, and thus enables us to reconstruct in their entirety those neurons that arborize in the neuropiles of both regions. Flies are held in a modified collar consisting of two halves held together on a magnetic base (Fig. 3). This assembly is used to dissect the dorsal and ventral brains attached, in two steps. First, the dorsal cuticle of the head and thorax is attached to one side of the collar with a tiny amount of cyanoacrylate glue (Loctite 404)(Fig. 3b-d). The exact amount is important and needs to be determined empirically. The proboscis and frons cuticle of the head are removed in a drop of saline (Olsen et al., 2007). Then the unglued side of the collar is removed and the attached side is turned into the horizontal plane and transferred into a Petri dish in a pool of saline (Wilson et al., 2005), and the legs removed (Supplementary Video 1). Next, the assemblage comprising the half collar attached to the partially dissected fly is transferred to primary fixative. Further steps are undertaken after 2h of fixation in 2.5% glutaraldehyde and 2.5% paraformaldehyde in 0.06M phosphate buffer. The second stage of dissection is undertaken in the same buffer. The head cuticle is removed, the collar turned 90° and the subesophageal ganglion and VNC dissected out. Even though the specimen is now fixed, the cervical connectives are structurally very weak after fixation and the specimen must be handled with great care to avoid fracturing its axons especially those of neurons that arborize in both ganglia. Despite these precautions, occasional dark profiles reflect the inevitable collateral damage of degenerating axons especially amongst afferent axons severed during the process of dissection.

### Speeding FIB-SEM: parallel imaging of hot-knife slices

Until now, FIB-SEM has been the slowest, and most costly imaging step in fly connectomics and could capture only small specimen volumes. For example, using FIB-SEM at an 8nm resolution the scan rate is only 1.25 MHz, covering a daily volume of just 25 x 10^3^ µm^3^ per day (Table 1, Method 1; Fig. 4a). Even though the dimensions over which a block can be milled, 400 x 300 µm (x, y) and ≤ 600 µm in z, could potentially include those of the fly’s entire brain, the area over which we could routinely collect a high-dose image stack using FIB-SEM without severe milling artifacts is far smaller than this (Xu et al., 2017). Meeting the need for increased ease and speed, during the last decade at Janelia we have developed successive generations of methods. In a first step, borrowed from an earlier precedent with light microscopy (McGee-Russell et al., 1990), we used the so-called hot-knife protocol to cut ultra-thick (∼20µm) slices (Hayworth et al., 2015; Fig. 5b) of an Epon (Poly/Bed 812, Ted pella) embedded brain coated with Durcupan. The choice of Durcupan was empirical, based on the superiority of this epoxy over Epon in having fewer streaks after FIB-SEM imaging (Xu et al., 2017). With the hot-knife method we could distribute the task of concurrent imaging amongst several slices, each imaged in a different machine. The female half-brain we report comprised about 13 20µm slices in a sagittal plane with a total imaged volume of up to ∼1.6 x 10^7^ µm^3^, and we stitched consecutive image stacks to yield a final volume (Hayworth et al., 2015). For the entire CNS the volume comprised parts of 27 25µm sagittal slices through the dorsal brain and 26 cross sections through the ventral nerve cord, much larger than the female half brain. Even using the hot-knife slicing strategem and imaging voxels at 8 nm, FIB-SEM typically covers not more than about 30 x 10^3^ µm^3^ per day per FIB-SEM machine (Table 1, Method 2; Fig. 4b), and thus initially was painfully slow. In parallel with hot-knife slicing and to increase FIB-SEM imaging speed yet further, we developed revised staining methods to a level that would enhance overall image contrast, and thereby support increased FIB-SEM imaging speeds. Our new staining method had to provide enhanced contrast optimally suited not only to detecting synapses but also simultaneously enhancing tissue and membrane contrast. For this we developed a heavy metal method to enhance staining, which has increased the imaging speed to 50 x 10^3^ µm^3^ per day per machine (Table 1, Method 3; Fig. 4c).

**Figure 4.**
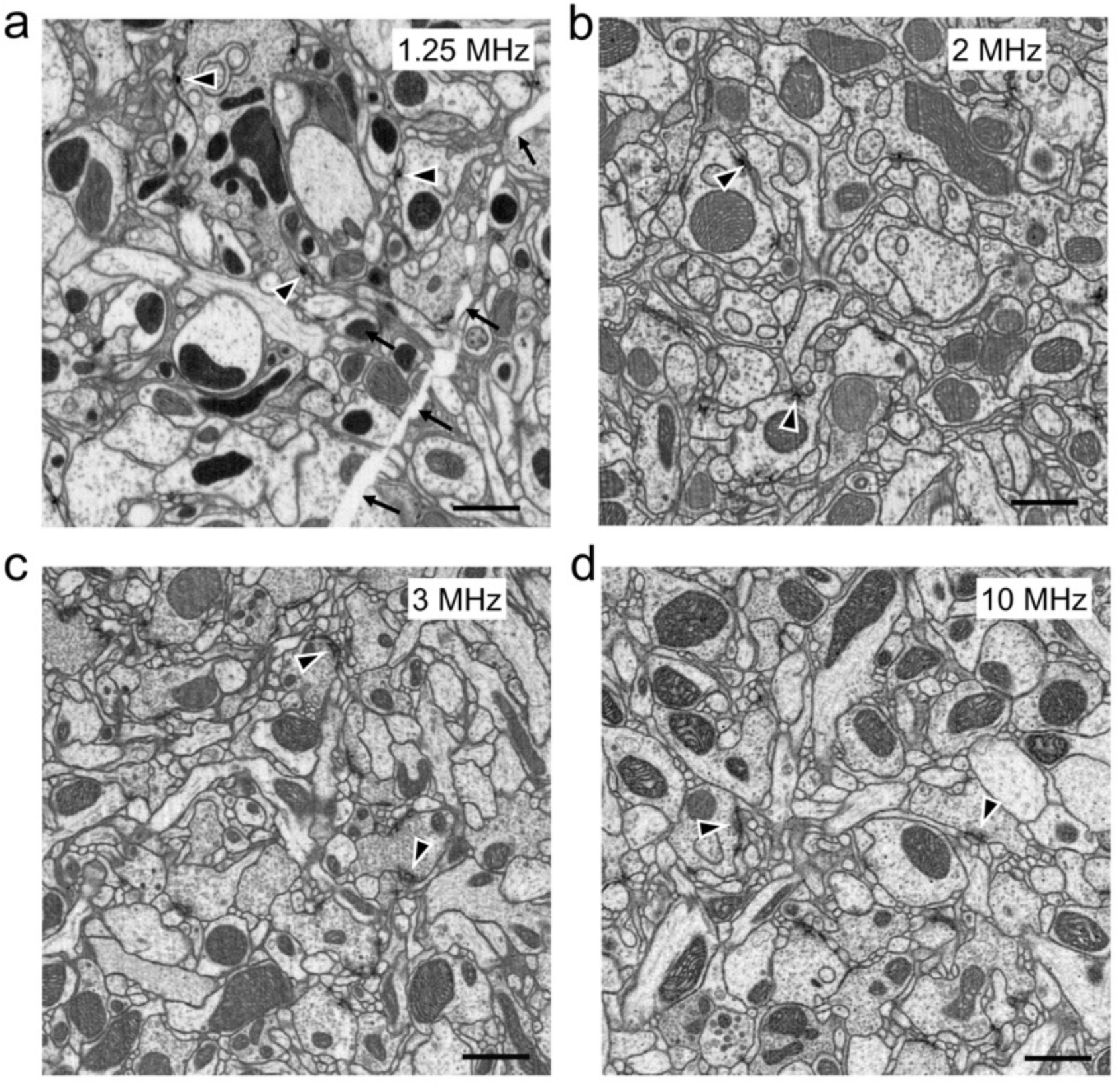
FIB-SEM images, from samples prepared with different methods at different scan rates. All produce images of comparable quality, collected at different imaging speeds. (a) High-pressure freezing with freeze substitution, HPF-FS to compare with (b) progressive lowering of temperature and low-temperature staining (PLT-LTS). (c) Heavy metal enhanced PLT-LTS method scanned at 1.5 times the rate (3MHz) as in (b). (d) PLT-LTS progressive heavy metal enhancement, imaged more rapidly than the preceding (c), according to Method 4 in Table 1. Synaptic profiles (arrowheads) are clear in all panels but unavoidable cracks (arrows in a) appear during HPF-FS specimen preparation. Scale bars: 1µm.

**Figure 5.**
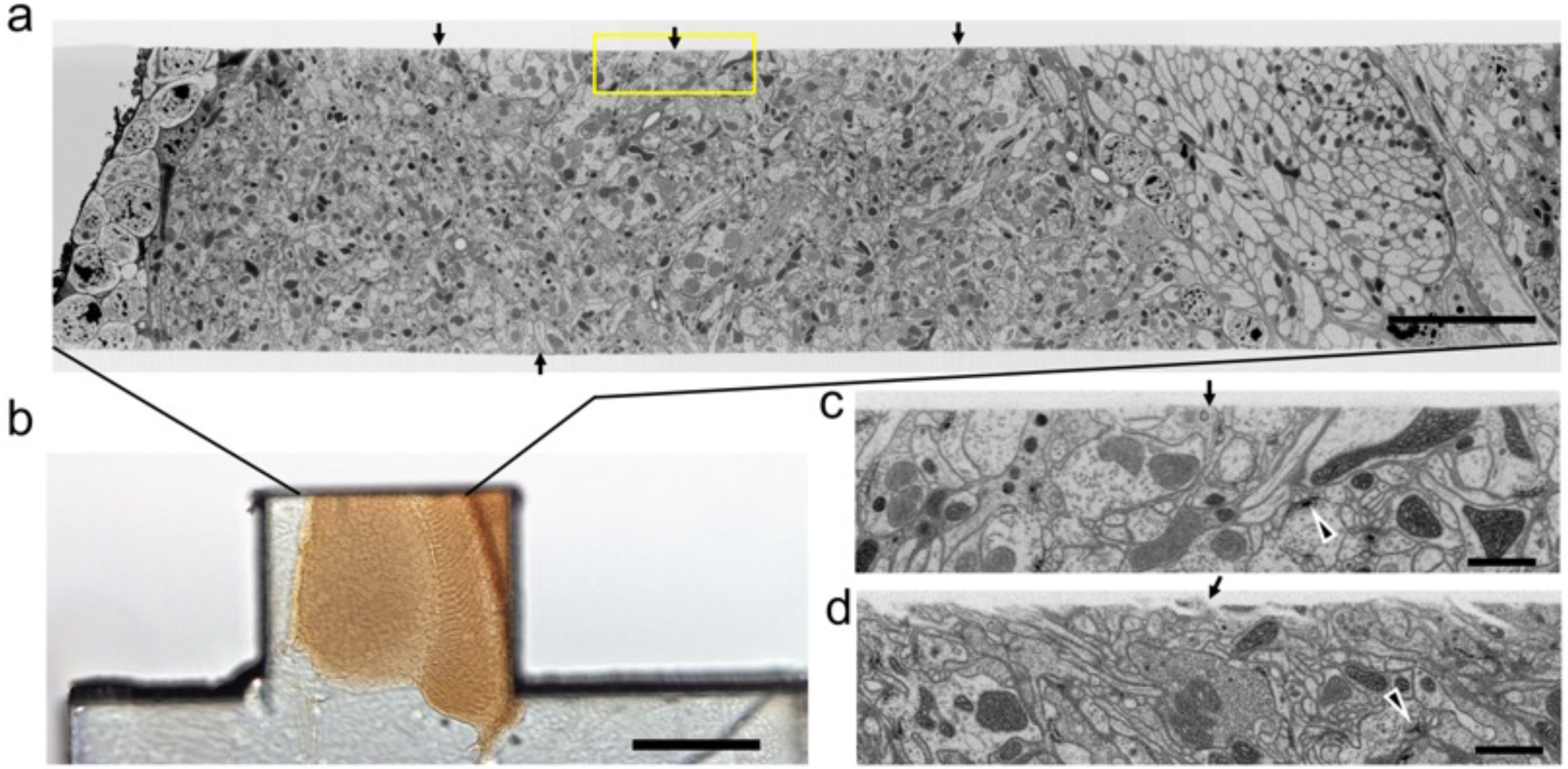
Quality of the cut surface of hot-knife slice. **a**. Entire FIB-SEM image of the medulla neuropile of a 20µm thick hot-knife slice shown in **b**. **b**. Light micrograph of 20µm slice of optic lobe. **c.** Enlarged area of one side of the cut surface, shown enclosed in yellow box in **a**. Arrows in **a** show the smoothness of the cut surface of the slice. Arrowhead in (c) and (d) shows selected synaptic profile. Arrows in (c) and (d) show the surface of the hot-knife slice, smooth (c) but occasionally rough (d). Scale bars: 10µm (a); 100µm (b); 1µm (c,d).

The increased contrast of these enabled us automatically to segment the profiles of neurons far faster than initially, while imaging at rates that now match those of ssEM (Xu et al., 2017, 2019). In consequence, imaging times are similarly reduced, thus saving in parallel the cost of expensive FIB-SEM imaging time. Moreover, although we stitched a few (up to 5) tiles per imaging plane, the isotropic image stacks so compiled did not require labor-intensive construction as montages, only that the image collected from each hot-knife slice be stitched to that of its neighbors in the stack.

### New *en bloc* staining methods

In our first attempts to image an entire fly’s brain using FIB-SEM we encountered successive problems, for which we developed a number of new methods (Table 1).

To obtain good brain preservation, especially with clear synaptic densities, we first used high-pressure freezing (HPF, Supplementary Fig. 2 and Table 1, Method 1) as a comprehensive method to analyze the entire *Drosophila* brain from successive slices. This method provided specimens with good morphology and image contrast (Takemura et al., 2015; Takemura et al., 2017; Shinomiya et al., 2019; Horne et al., 2019), close to the living state, but brain blocks developed cracks and lacked the cutting properties for the hot-knife protocol required to sample large volumes of the fly’s brain. We therefore switched to progressive lowering of temperature and low temperature staining (PLT-LTS) to image the brain (Figs. 2c, 5a; Table 1, Method 2; Supplementary Table 1). This preserved all the favourable features of high pressure freezing, and cured the problem of cracked blocks, staining T-bars at synaptic sites dark, as well as enabling us to slice the brain using the hot-knife method (Hayworth et al., 2015). Nevertheless the overall image speed was still slow. For example, using the PLT protocol above, FIB imaging took ∼80 days per slice, but without incurring specimen cracks, while retaining good hot-knife cutting properties to allow simultaneous parallel imaging of multiple slices. To improve imaging speed we also increased tissue contrast by means of supplementary heavy metal enhancement, using potassium ferricyanide and lead aspartate (Table 1, Method 3). In addition, we used BSA coating (Fig. 6a) to offset any disadvantageous (as shown in Fig 5d) effects of different mechanical properties on the cutting process as well as provide structural features that allow SEM focus optimization prior to tissue imaging.

**Figure 6.**
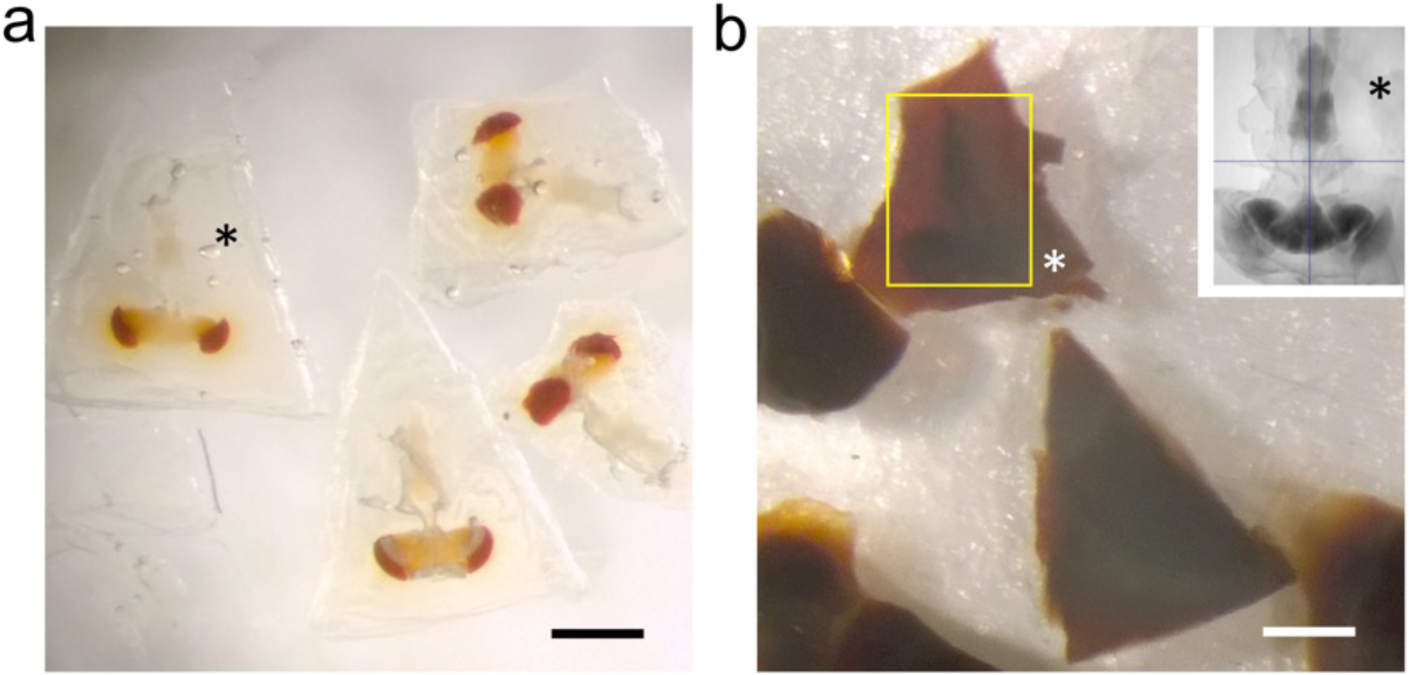
BSA coating of dissected brain. **a**. Fixed specimens in trapezoidal blocks after coating with BSA. **b**. Osmication and staining with metal salts darkens the BSA, but leaves the brain’s outline still visible (yellow rectangle). Different zones in the surrounding BSA reveal successive additions of the cross-linked protein. Inset: X-ray tomogram of a single CNS. Asterisks in all images show the BSA coat in different stages of processing. All scale bars: 500µm.

However, the heavy metal exacerbates the poor cutting property of the brains for thick sectioning. We overcame this problem by first coating the brain with bovine serum albumin (BSA, Fig. 6a). This provides a smooth surface to the cut edge (Fig. 5a and 5c) that was useful later in improving the registration of cut surfaces between consecutive slices, as well revealing structural features that provide SEM focus to be optimized before the tissue itself is milled. This method proved superior to that of Hildebrand et al. (2017) in diminishing the gap between BSA and the fly brain tissue, and in providing a more extensive surface of contact. However, a countering disadvantage was that the BSA darkens and obscures the outline of the enclosed fly brain (Fig. 6b). To overcome this problem, we imaged the tissue with X-ray tomography (microCT) prior to FIB-SEM, a combination of methods that enabled us to select the desired brain region with great accuracy.

To obtain this improved staining, fly heads were dissected and prepared in a metal collar, as given above (Fig. 3), pre-fixed in a mixture of 2.5% of each of glutaraldehyde and paraformaldehyde at room temperature (22°C) for 2 h, and then post-fixed in 0.5% OsO_4_ for 40min at 4°C, followed by three 10-min washes in water; then heavy metal enhancement in 0.8% K ferrocyanide for 2h at 0°C and 0.5% uranyl acetate, 30min at 0°C; a wash, and then staining in lead aspartate overnight at 4°C; followed by a distilled water wash, and then 20min in 0.8% OsO4 at 0°C. Dehydration is followed by embedment in Epon or Durcupan.

In addition to applying this protocol to adult flies, we also developed a method by combining ferrocyanide reduced osmium-thiocarbohydrazide-osmium (R-OTO: Willingham and Rutherford, 1984) to PLT-LTS that enabled us to image the brain of first-instar larval *Drosophila*. This is is smaller and thus unlike the adult brain did not require hot-knife slicing. This progressive heavy metal enhancement method (Table 1, Method 4, Fig. 4d) applied to both larva and adult used the advantages of Method 3 while increasing tissue contrast; and in combination with the brain’s small size it enabled us to collect an image stack with a high FIB-SEM imaging scan rate of 10MHz to achieve 200 x 10^3^ µm^3^ per day per machine at 8nm resolution (Table 1, Method 4) 8 times faster than Method 1.

### X-ray tomography

To locate regions of interest, we routinely employ X-ray tomography of osmicated specimens viewed *en bloc*, using an Xradia Versa 3D XRM-510 to preview the specimen and select out those specimens having cracks, vacuoles or distortions that would have wasted valuable imaging time on flawed specimens. This important step also enables us to identify the coordinates of imaged structures prior to trimming the block to a specific depth for FIB milling (Takemura et al., 2017), in a combination of methods that enabled us to select the desired brain region with great accuracy. Both selections, of the region of interest and its depth, offer considerable prospective savings against wasting time to mill and image through unwanted sample areas, during the lengthiest but most valuable step in the entire process. Executing this step requires some experience however. Scrupulous preservation and integrity of the brain is required because of the large time investment in fixation and FIB-SEM imaging made after the initial dissection, and because superior fixation can only be selected at the end of these steps, after a lengthy period of imaging that would otherwise be wasted on an inferior sample.

## Conclusions

Dissection and fixation are the first essential steps to view cells and organelles. In previous studies, dissection of the *Drosophila* brain has generally been minimal, involving only removal of the eye and lamina of each side, and fixation is aided by the brain’s tiny dimensions, <150µm along the head’s anteroposterior axis (Peng et al., 2011), and hence well suited to EM. Most conventional primary fixation methods employ aldehydes, especially primary fixation by the formaldehyde/glutaraldehyde (PFA/GA) mixture with high osmolarity introduced by Karnovsky (1965) >50 years ago. The advantage of this and other double-aldehyde fixatives is that they provide a universal method that needs no refinement for particular nervous systems, even if many simple invertebrate nervous systems do not in fact fix well with it. *Drosophila* is generally well preserved with aldehyde fixation methods (Wolff, 2011; Meinertzhagen, 1996), but for neuropiles a general problem is to capture neurites as profiles that are round in cross section and well separated from those of their neighbours, well suited to automated segmentation. Most EM using previous techniques preserves many neurites only as flattened and polymorphic profiles, however, a usual condition in published EM images, and makes the continuity of these profiles hard to follow in an image stack. To enhance membrane density, high-pressure freezing and freeze substitution (HPF-FS: Walther and Ziegler, 2002), and ferrocyanide reduced osmium-thiocarbohydrazide-osmium ligand binding (R-OTO: Willingham and Rutherford, 1984), have all been used, but each has its own shortcomings particularly for intact insect brain tissue.

Addressing these deficiencies, we report a number of methods adapted to the analysis of synaptic circuits in the *Drosophila* brain (Supplementary information). The detailed protocol we present for *Drosophila* incorporates various component methods which, in differing combinations, are likely to suit the fixation of brains in other model species. Individual steps in our protocols, for example BSA coating for hot-knife slicing of entire brains and our dissection protocols, are for example equally applicable to the connectomics of other species. They are the product of a decade of our development from earlier protocols. Each offers particular advantages, but most important for our purposes, we report a method to improve the imaging speed of FIB-SEM by adopting novel ways to increase specimen contrast, and we apply these to an entire microdissected hot-knife sliced fly’s brain comprising connected sub- and supraesophageal ganglia. Our methods are adapted to a FIB-SEM imaging mode and reliably recover fixed neurons as round cross-sections, suitable for machine segmentation (Parag et al., 2015), with dark synaptic profiles suitable for automated synapse detection (Huang et al., 2018). The numbers of the latter match closely the numbers of those identified by human proof-readers (Shinomiya et al., 2019) and so are considered accurate.

In aggregate our collective methods, those reported here and others developed at Janelia (Supplementary information Methods), provide a means for semi-automatic segmentation of *Drosophila* neurons and automated synapse detection. In particular, our staining methods now provide an excellent compromise between specimen contrast and accelerated FIB-SEM sectioning speed. Imaging speed may be further enhanced using higher specimen contrast to yield usable images yet more quickly, however; and in the future also possibly by using gas cluster milling (Hayworth et al., 2019) combined with SEM with multi-beam imaging (Eberle and Zeidler, 2018). Even so, many sensory inputs to both brain regions are necessarily removed when their axons are severed, and these leave behind degenerating afferent axons, which yield electron-dense profiles. Darkened degenerating axons visible in EM are known to appear with a very rapid onset (Brandstätter et al., 1991) and in our case are thought to signify those axons that were severed, or also possibly simply stretched, during the relatively short period of dissection and immediate fixation.

The rationale for our PLT-LTS method is based on previously reported size measurements in cells prepared for EM. PLT-LTS gives the tissue intense staining and fewer structural alterations than routine dehydration and *en bloc* staining. Using a lower concentration of ethanol (<70%) during dehydration causes the tissue to swell whereas with dehydration in absolute ethanol tissue the tissue shrinks (Konwiński et al., 1974). Dehydration at low temperatures can minimize these size and shape changes. We also found that after staining tissue at 0-25°C in acetone- or ethanol-based uranyl acetate after routine fixation, the FIB-SEM images showed improved contrast compared with routine staining with aqueous UA at 4°C (Supplementary Fig. 3). Using acetone gave the best results in tissue contrast but the hot-knife cutting properties were worsened, making a compromise necessary. The PLT-LTS method helps to provide uniform osmication and staining, with less chance of distorting the fine structure. The method works well on the entire adult *Drosophila* brain as well as that of the first-instar larva. To extend the PLT technique this protocol could be further improved by introducing lead acetate, tannic acid, imidazole, phosphotungstic and organic solvent soluble stains into the protocol.

Finally, our method incorporates an important advance in being able to preserve both parts of the CNS intact while these still remain connected, and thus make it possible to image pathways between the supraesophageal and subesophageal ganglia of the brain and the cells that arborize in both. Preserving the continuity of pathways through the connectives ensures retention of the integrity of descending inputs to the many lineages of subesophageal neurons (Shepherd et al., 2019), as well as complementary ascending pathways. Only by retaining both halves of the brain can cells with neurite arbors in both be preserved complete, which as far as we are aware has not previously been reported for EM. An unavoidable consequence of removing the brain from the fly’s head is, even so, that many sensory inputs to both brain regions are necessarily removed when their axons are cut, and these leave behind degenerating afferent axons, yielding electron-dense profiles. These we regard as the small inevitable price to pay for the opportunity our methods provide to identify the brain circuits formed by the majority of intact well-preserved axons.

## Acknowledgements

We thank the FlyEM team at the Janelia Campus of HHMI for assistance, HHMI for support, and Dr. Yalin Wang and Weiping Li in particular for valuable discussions.

## AUTHOR CONTRIBUTIONS

Z. L. undertook all fixations and EM analyses, prepared all figures, and helped prepare the manuscript; C.S.X. and K.J.H. undertook all FIB imaging and image alignments; C.S.X., K.J.H., P.R. and Z.L. undertook EM analyses; P.R., S.M.P, L.S, H.F.H. supervised the project; and G.M.R. and I.A.M. prepared the manuscript with input from all coauthors.

## Supplementary information

**Supplementary Fig. 1.**
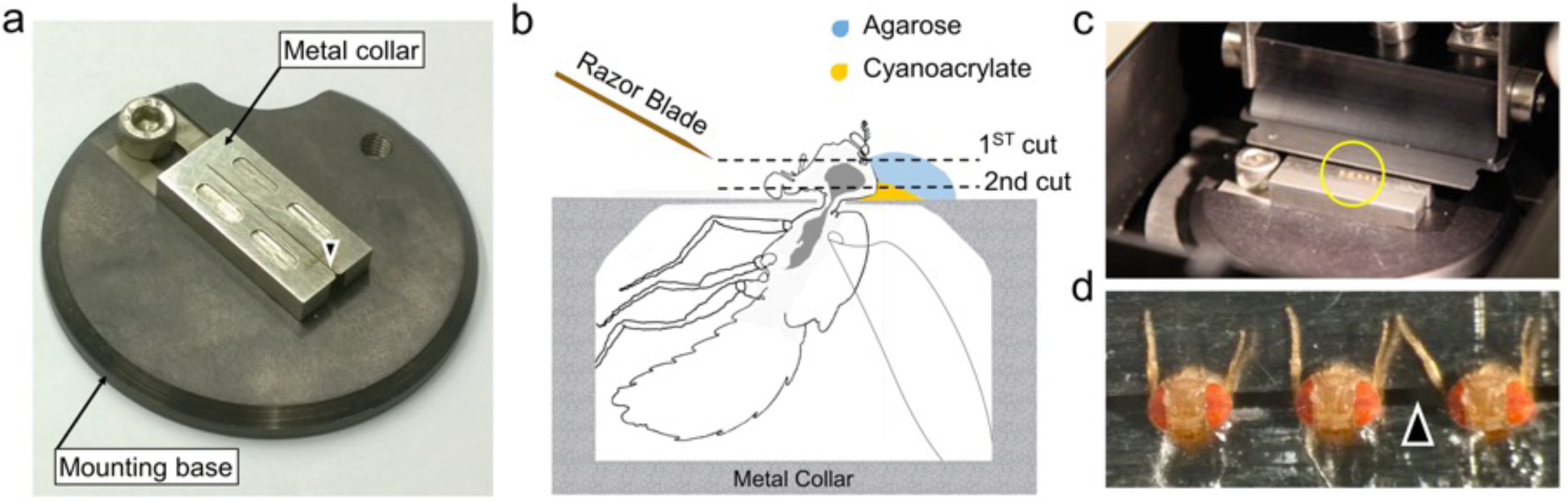
Dissection collar for *Drosophila* Vibratome slice. a: Custom-made dissection collar mounted on a Vibratome sample loading base. A slot holds a row of flies captive at their necks, held on a thin metal shim between four reservoirs for saline. b: Side view of a fly in the collar with its head protruding through the slot, mounted with cyanoacrylate cement (Loctite) and covered in 5% agarose to stabilize the entire head during slicing, when cut at an angle by a thin high-carbon steel Feather double-edge razor blade. The first cut removes the anterior head cuticle, after which a single drop of fixative starts fixation, immediately followed by a second cut, which removes a 200µm slice. c: Vibratome slice of five fly heads (circle) in a collar. d: Enlarged frontal view of three of the heads in the collar (from c).

**Supplementary Fig. 2.**
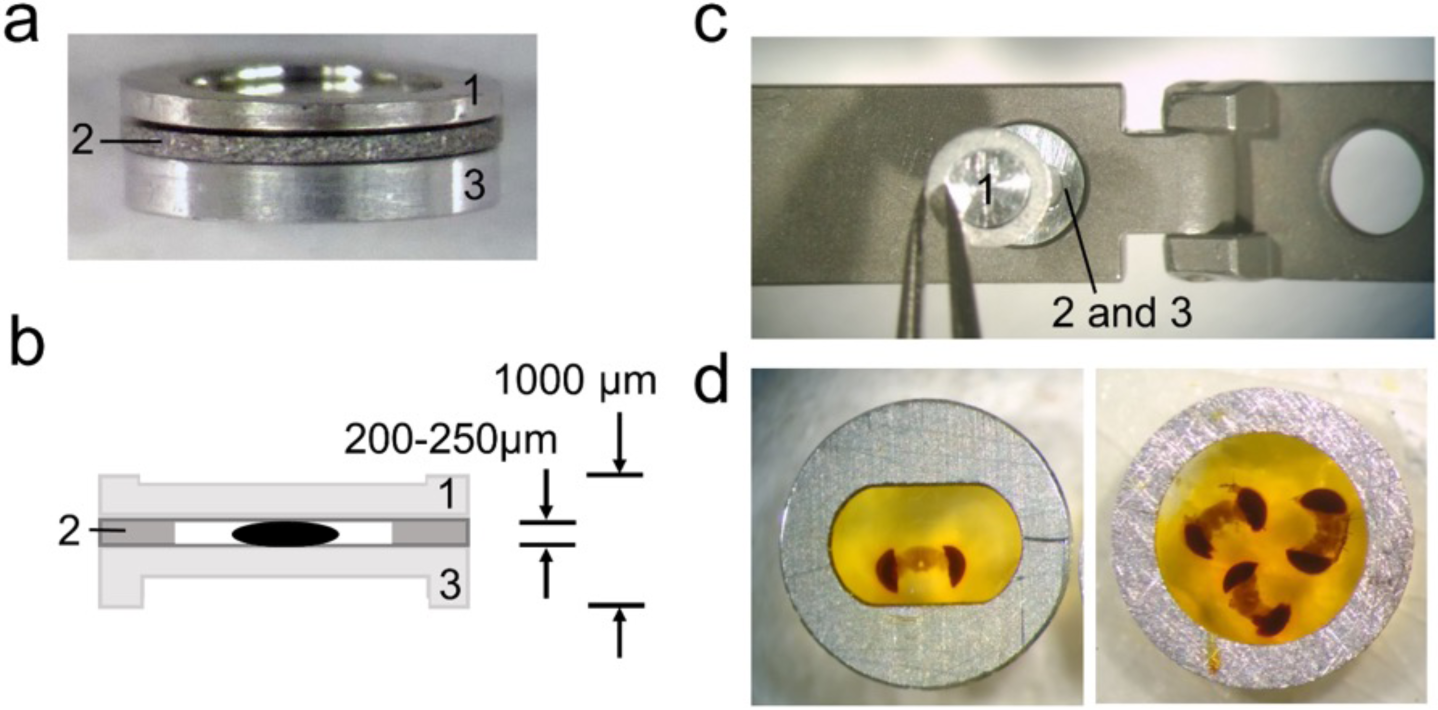
High-pressure carrier for freezing *Drosophila* brains. a: Custom-made aluminum sandwich carrier for high-pressure freezing fly heads, comprising a machined annulus (2) sandwiched between two annular plates (1,3). b: Cross section of sandwich in a, with 200-250 µm Vibratome slice of a fly head (black profile) in specimen annulus (2) supported between the two annular plates (1,3) coated on their inner faces with lecithin. c: Final assembled thickness is 1000 µm to fit in the specimen holder of a Wohlwend HPF Compact 01 High-Pressure Freezing Machine (Wohlwend GmbH, Sennwald, Germany). d: Samples in specimen annulus (2 in a-c) after polymerization. The hinged top and bottom layers (1,3 in a-c) are removed before freeze substitution. During freeze substitution the medium can substitute from both free surfaces. Specimens are surrounded by filler (20% BSA filler in water), yellow after polymerization. A specimen annulus having a round well provides a larger area for freeze substitution than one with an elliptical well. Specimens are easily removed from the annulus by cutting the latter along one diameter with a single-edge razor blade.

**Supplementary Fig. 3.**
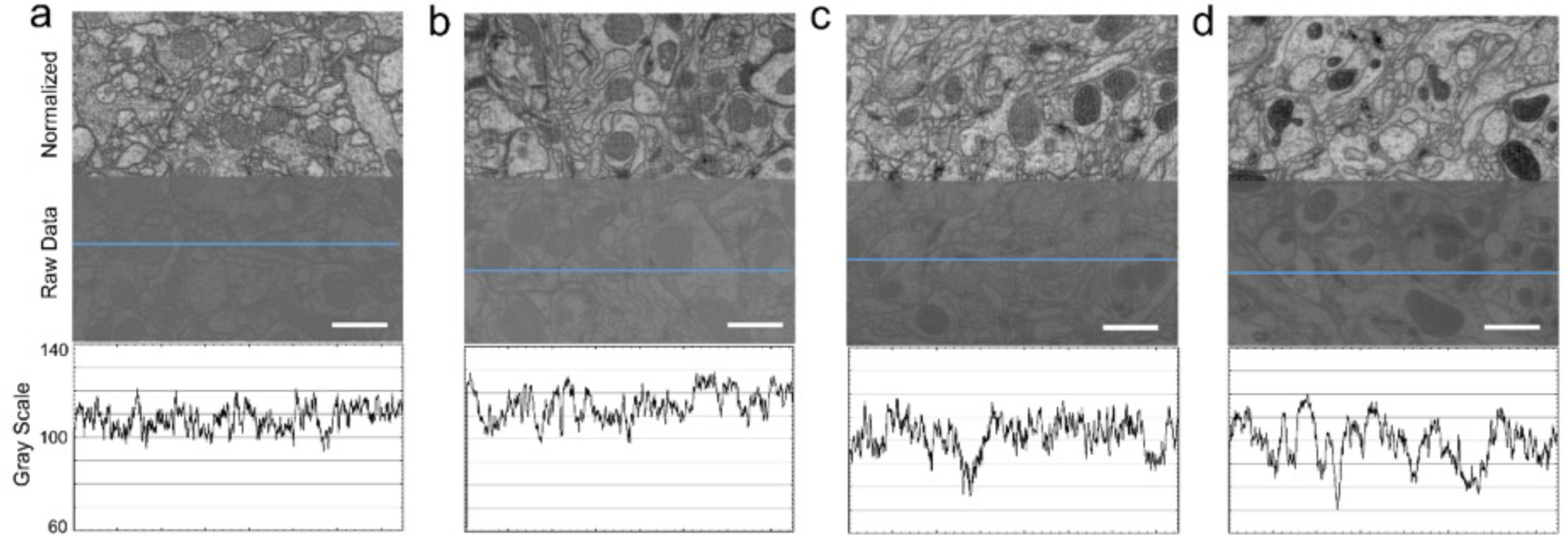
Comparison of the contrast in FIB-SEM images using UA staining. Comparison between *en bloc* staining of aqueous UA (a, b) and organic-solvent based UA (c, d) on adult *Drosophila* brains. There are three parts to each figure. The top part is the normalized half of the raw image; the middle part is the raw data without changing the range of pixel intensity values; the bottom part shows the “Plot Profile” (Fiji - ImageJ) required to display a 2D graph of the pixel intensities along the blue line within each image. The range indicates the contrast, wider fluctuations reflecting higher contrasts. (a) shows the tissue *en bloc* staining with aqueous 0.3% UA; (b) shows tissue after 1% UA overnight at 4°C with a conventional fixation and dehydration procedure. (c) shows the tissue staining with 0.3% UA in ethanol; and (d) shows the staining with 0.2% UA in acetone using the PLT-LTS procedure (see Supplementary Table 1). The overall contrast produced by aqueous UA staining is lower than the staining contrast with UA in ethanol or acetone. Using acetone gave the best results in tissue contrast. Scale bars: 1 µm.

**Supplementary Table 1:**
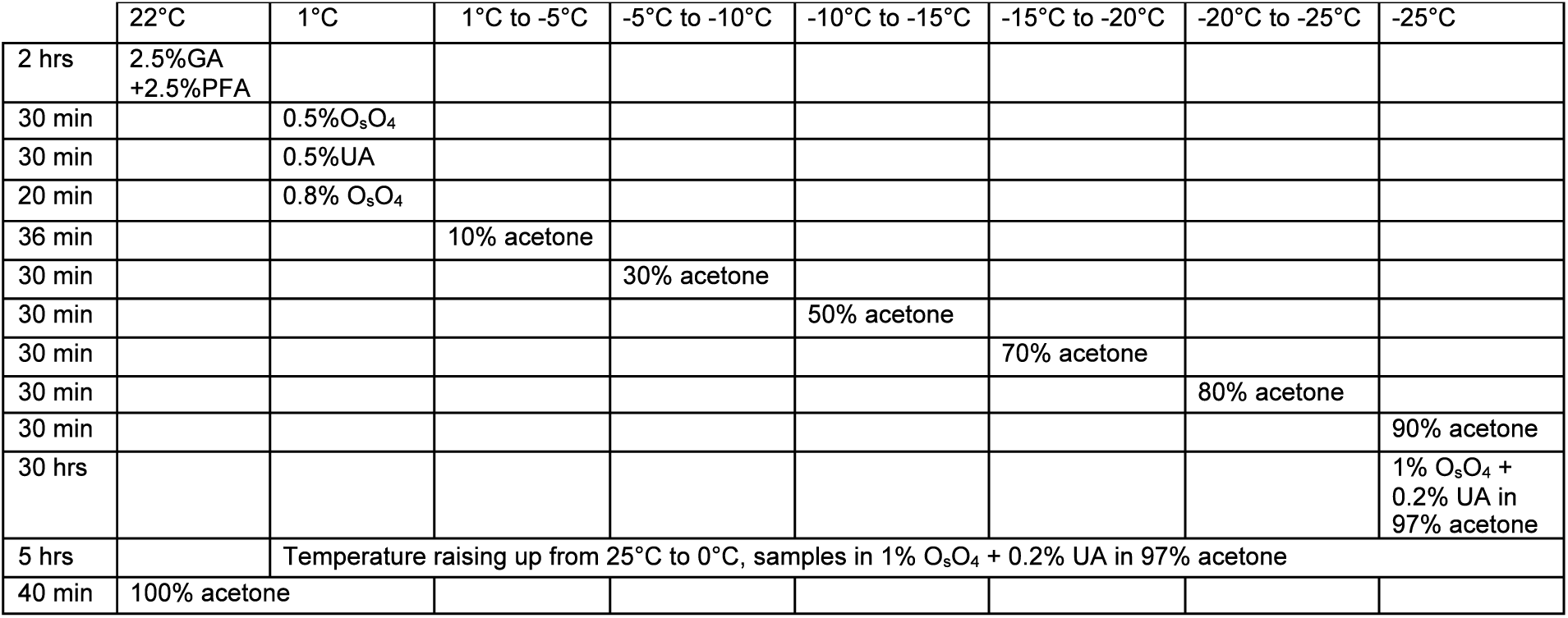
Timetable of temperature and solvents changing of PLT-LS processing

**Legend to Video 1**. Steps in the dissection of the *Drosophila* CNS (see Fig. 3). First, loading flies into a dissection collar (Fig. 3c) and gluing the head and thorax onto the collar (Figs. 3c, 3d). Next, dissection steps 1, 2 and 3 (Figs. 3b, 3e, 3f). Next: final step to dissect out the CNS after primary fixation (Fig. 3g). Chief dissection tools used: custom made metal dissection collar; fine forceps with sharpened tips; Feather carbon steel razor blades; blade holders and breakers with concave-convex jaws; cyanoacrylate glue (Loctite 404).

**Legend to Video 2**. High-resolution 4-nm per pixel FIB-SEM video of the protocerebral bridge showing the quality of PLT-LTS sample preservation in the image stack containing the single image presented in Fig. 2c.

## Methods

### Animals and main steps

As specimens we used Canton-S G1 x w^1118^ wild-type ∼5-day adult *Drosophila melanogaster* maintained at 23-25 °C on standard fruit fly medium. To prepare *Drosophila* brain tissue specifically to image the entire *Drosophila* brain by FIB-SEM we developed a number of general methods (Table 1), each offering an improvement over the previous one, and we report only our final method in the Results, even though previous methods provide alternative advantages for different aims. We used conventional primary fixation according to the protocol of Takemura et al. (2013) for ssEM and modified this in one of three ways to enable us to minimize the time required for FIB-SEM of an entire *Drosophila* brain. Chief among these, we adopted the hot-knife method (Hayworth et al., 2015) to view several such volumes from successive 20µm slices imaged in parallel in different machines, and subsequently stitched these to generate a single volume. Each slice comprised 2500 8-nm FIB-SEM images.

### Method 1). HPS-FS

In the first modification we applied *High Pressure Freezing (HPF) after primary fixation*. The fly’s fixed brain was sliced in a custom-made dissection collar (Supplementary Fig. 1a) mounted on the slicing base of a Vibratome. We cut 200µm slices using a Leica Vt1000 Vibratome (Supplementary Fig. 1c); the slices were fixed in 2.5%GA + 2.5% PFA for 10-15 min, transferred to 25% aqueous bovine serum albumin for a few minutes, and then loaded into a 220µm deep specimen carrier sandwich, and high-pressure frozen in a Wohlwend HPF Compact 01 High Pressure Freezing Machine (Wöhlwend Gmbh, Sennwald, Switzerland). This arrangement of specimen carrier sandwich (Supplementary Fig. 2a-c) was chosen instead of a two-hat carrier, widely used in the field for large samples (Murk et al., 2003; McDonald, 2009). After freeze substitution (FS), slices (Figs. 2d,e) were embedded in Durcupan (ACM Fluka) epoxy resin (Xu et al., 2017; Horne et al., 2019; Tanimura et al., 2018; Shinomiya et al., 2017, 2019), in preparation for FIB-SEM (Knott et al., 2008; Xu et al., 2017). The choice of Durcupan is empirical, based on the superiority of this epoxy to Epon in having fewer streaks after imaging (Xu et al., 2017). On the other hand, HPF-FS samples do not cut well for ssEM or during trimming, and to avoid its use we therefore mostly discontinued this freezing method and developed a method for chemical fixation using dehydration by progressive lowering of temperature (Hayworth et al., 2015), see Method 2.

### Method 2) PLT-LTS

The fly brain was dissected out by using a metal dissection collar (see Fig. 3), then given primary fixation in 2.5%GA + 2.5%PFA in 0.06M PB for 2 hours at 22°C, then washed 3×10 min in 0.06M PB followed by cacodylate buffer. Specimens were next exposed for 30min to 0.5% osmium tetroxide in 0.05M cacodylate buffer. Then the following procedure was adopted using a protocol we have reported previously in which brains are fixed chemically and processed using dehydration by *progressive lowering of temperature* (PLT)(**Supplementary Table 1**) also referred to as C-PLT (Hayworth et al., 2015), which reveals synapses having high-contrast organelles. In our current method for adult *Drosophila*, we changed the buffer from 0.1M to 0.06M, which we have found decreased the electron density of the cytoplasm. We found this decrease by examining multiple specimens, and despite some individual variation between these.

### Method 3) PLT-LTS heavy metal enhancement

In addition to PLT we have employed *heavy metal contrast enhancement*, an improved protocol for *Drosophila* brains that yields an excellent compromise between optimal contrast, sectioning speed and morphological preservation for FIB-SEM, and is that is also compatible with hot-knife slicing. This method yields high overall electron contrast for membranes and other cellular structures, but a relatively lower contrast for synapses. After dissection (see Fig. 3) and primary fixation as in Method 2, we could either coat the dissected CNS with BSA in order to undertake hot-knife slicing, or without coating, and then wash the specimens for 3×10 min in PB and then cacodylate buffer, post-fix them in 0.5% osmium tetroxide in 0.05M sodium cacodylate buffer, and finally treat them with 0.8% potassium ferricyanide in buffer for 2 h at 4°C. After washing in water, we incubated the tissue in 0.5% aqueous uranyl acetate (UA) for 30 min at 4°C followed by *en bloc* staining in lead aspartate at 4°C overnight, or for 4 hours at 22°C, and then after further washing in water, for 20min in 0.8% OsO_4_. For PLT, we placed specimens in a Leica AFS freeze-substitution chamber and dropped the temperature from 4°C to −25°C, and increased the concentration of acetone or ethanol for 20 min in each of 10%, 30%, 50%, 70%, 80%, 90% and 97% (Supplementary Table 1). *En bloc staining* and further osmication used a cocktail of 1% osmium tetroxide and 0.2% UA in 97% acetone or ethanol at −25°C for approximately 30 h, warming to 22°C for final dehydration, then infiltration in acetone or propylene oxide with Epon or Durcupan (Xu et al., 2017; Hayworth et al., 2015). This protocol is current and has been used to analyse the connectome of half a female fly’s brain. For method 3 we did not use OTO because this resulted in preparations with inferior cutting properties.

To image the entire CNS using FIB, we first needed to cut the preparation into 20-30µm slices that could be individually handled. For this we improved the hot knife cutting properties using a custom-made ultrathick sectioning microtome (Hayworth et al., 2015). To improve the cutting properties of the brain and preserve the integrity of its external surface, which is easily distorted, we developed a new method, enclosing the brain in a 25% solution of bovine serum albumin (BSA) in 0.06M phosphate buffer (PB) after primary fixation. This process relies on cross-linking the BSA after aldehyde fixation. We do this by placing three drops in a Petri dish, the first containing 25% BSA, the second containing fixative, and the third buffer wash (PB). The brain is first transferred from the BSA (drop 1) to the fixative drop (drop 2) to coat it with a thin layer of fixed protein, and next transferred to the buffer wash drop (drop 3), then back to the BSA drop. This sequence was sometimes repeated, to ensure that a thin layer of BSA adhered to the ventral surface of the specimen. Using this sequence, we added a drop of fixative on top of the BSA droplet containing the sample, and waited until the BSA polymerized. We then cut the polymerized BSA into a regular trapezoid containing the orientated sample at its center (Fig. 6a). The inclusion is then carefully lifted and transferred to a droplet of buffer wash. After osmication, heavy metal staining and further processing, the BSA coating layer darkened (Fig. 6b). We used X-ray tomography to provide a means to view the sample’s profile and orientation from its opaque BSA coating (Fig. 1).

### Method 4) PLT-LTS progressive heavy metal enhancement

For contrast, we used OTO to provide high contrast, because we were not concerned about the quality of cutting. We used this method to image the larval brain, which is too small to require hot-knife subdivision. After tissue dissection and pre-fixation as in methods 2 and 3, we osmicated tissue in 1% OsO_4_. This was followed without washing by 1.5% K ferrocyanide, then a complete wash and finally, a transfer for 15 min to 1% thiocarbohydrazide at 22°C, followed in turn by a complete wash then 2% osmium for 30 min at 22°C. After osmication we stained in lead aspartate for 30 min at 55°C, followed by 1 h at 22°C. The tissue was then dehydrated using the PLT method as in Method 3, followed by low temperature *en bloc* staining in either 0.2% uranyl acetate in acetone, or 1% EPTA in 97% ethanol. Specimens were infiltrated and embedded as for methods 1-3 in Epon or, in the case of FIB, Durcupan.

